# Design, Fabrication, and *Ex-vivo* Validation of an Active Capsule Endoscope

**DOI:** 10.64898/2026.06.14.732152

**Authors:** Deepak Kumar Dinkar, M Hasan Shaheed, Kaspar Althoefer, Mohamed Adhnan Thaha

## Abstract

**Background and Aims:** Active capsule endoscopy could advance gastrointestinal diagnostics by enabling controlled navigation beyond passive peristalsis. However, current systems are often limited by inefficient propulsion, high power demands, or reliance on external actuation. Herein, we designed, developed and evaluated a novel electromagnetic impact-actuated capsule endoscope incorporating a ferromagnetic rail-enhanced locomotion mechanism.

**Methods:** The capsule employed an internal electromagnetic actuator comprising a movable coil-armature assembly guided along a ferromagnetic rail and surrounded by permanent magnets. Controlled current pulses generated reciprocating motion and propulsion through momentum transfer. Bench-top testing using a deformable intestinal model assessed locomotion and power consumption. *Ex-vivo* experiments were subsequently performed in porcine intestine under dry and physiologically simulated wet conditions. Transit speed, power consumption, and system stability were recorded.

**Results:** Bench-top testing demonstrated stable propulsion at speeds up to 8.5 mm/s with a mean power consumption of 84 mW. During *ex-vivo* evaluation, mean capsule velocities were 1.95 mm/s and 7.2 mm/s under dry and wet conditions, respectively. Average power consumption was 96 mW and 193 mW. The actuator maintained reliable locomotion while preserving a compact system volume of ∼6.19 cm^3^. Lubricated conditions, representative of the intestinal environment, resulted in enhanced propulsion efficiency despite a concomitant increase in instantaneous power consumption.

**Conclusion:** The electromagnetic impact-actuated capsule demonstrated reliable locomotion in biologically relevant *ex-vivo* environments while maintaining compact dimensions and moderate power requirements. Ferromagnetic rail-enhanced flux concentration offers a promising propulsion strategy for future actively navigated and therapeutic capsule endoscopy platforms.

## 1. Introduction

Since its first clinical application in 2000, capsule endoscopy (CE) has revolutionised gastrointestinal (GI) imaging by enabling minimally invasive visualisation of the digestive tract [1-3]. The technology has proven particularly valuable for examination of the small intestine, a region that remains difficult to access using conventional flexible endoscopy because of its length, tortuous anatomy, and relative inaccessibility [4]. Over the past two decades, CE has become an established diagnostic modality for the investigation of obscure gastrointestinal bleeding, Crohn’s disease, iron deficiency anaemia, small bowel pathology, and colorectal neoplasia [5-8]. Advances in image acquisition, wireless communication, localisation technologies, and artificial intelligence-assisted interpretation have further expanded its power management, diagnostic capabilities, increasing interest in its application for colonic evaluation and bowel cancer screening [9-12]. Owing to its non-invasive nature, high patient acceptability, and ability to visualise the entire gastrointestinal tract during a single ambulatory examination, CE has emerged as an attractive alternative to conventional endoscopic procedures.

Despite these advantages, commercially available capsule endoscopes remain fundamentally limited by their reliance on passive propulsion through natural peristalsis and gravity. As a consequence, capsule transit is largely uncontrolled, resulting in variable examination times, incomplete visualisation, and inconsistent diagnostic yield [13]. Incomplete examinations have been reported in up to 20% of studies, particularly when delayed gastric emptying or prolonged intestinal transit occurs [14]. Furthermore, the inability to actively control capsule position, orientation, or velocity prevents targeted inspection of suspicious lesions and limits the development of therapeutic capabilities such as biopsy acquisition, localised drug delivery, and tissue intervention. Battery capacity remains another significant constraint, requiring a compromise between image quality, frame rate, wireless communication, and operational duration [15-18].

To overcome these limitations, substantial research efforts have focused on the development of actively propelled capsule endoscopes capable of controlled locomotion and navigation. Various propulsion strategies have been proposed, including externally actuated magnetic systems, legged mechanisms, inchworm-inspired robots, paddling devices, screw-driven propulsion systems, and hybrid locomotion architectures [19-24]. While many of these approaches have demonstrated proof-of-concept functionality, they frequently require bulky external magnetic platforms, complex mechanical assemblies, high power consumption, or sophisticated control algorithms that complicate miniaturisation and clinical translation. Achieving a balance between propulsion efficiency, device size, mechanical simplicity, and energy consumption therefore remains one of the principal challenges in active capsule endoscopy research.

Among the emerging propulsion concepts, impact-based locomotion systems offer a promising alternative because they can generate substantial propulsion forces using relatively simple internal actuation mechanisms. By accelerating an internal mass and transferring momentum to the capsule body through controlled impact, locomotion can be achieved without the need for external magnetic guidance or complex appendages. However, the efficiency of such systems is strongly dependent upon the electromagnetic force available for accelerating the moving mass. Improving force generation while maintaining compact dimensions and low power consumption remains a significant engineering challenge.

To address this limitation, we developed a novel electromagnetic impact-actuated locomotion system incorporating a ferromagnetic rail-enhanced magnetic circuit [23]. The system employs a movable coil-armature assembly travelling along a ferromagnetic rail positioned within a circumferential permanent magnet array. In addition to guiding armature motion, the rail concentrates magnetic flux through the actuator region, thereby increasing electromagnetic force generation and improving propulsion efficiency. This architecture aims to provide reliable capsule locomotion while maintaining a compact form factor and moderate power requirements suitable for future clinical translation.

In this study, we present the bench-top and *ex-vivo* evaluation of this active capsule endoscope. Bench-top experiments were performed to characterise actuator performance, propulsion behaviour, and energy consumption under controlled mechanical conditions. Building upon these findings, *ex-vivo* trials were conducted using porcine intestinal segments under both dry and physiologically simulated wet conditions to investigate locomotion efficiency, power consumption, and environmental influences on propulsion performance. By systematically evaluating the device across increasingly realistic test environments, this work provides critical insight into the feasibility of rail-enhanced electromagnetic locomotion and establishes an experimental foundation for future *in-vivo* validation and clinical development of actively navigated capsule endoscopy systems.

## 2. Experimental Methods

### 2.1 Electromagnetic Locomotion System Design

The active capsule endoscope was based on a novel electromagnetic impact-actuated locomotion mechanism developed for controlled movement within the gastrointestinal tract. The propulsion system consisted of a cylindrical housing incorporating a circumferential array of permanent neodymium magnets, a ferromagnetic central rail, and a movable coil-armature assembly, as shown in Fig. 1. The permanent magnets were arranged with identical pole orientations facing inward to generate a concentrated magnetic field directed towards the central rail. The movable armature comprised a steel sled wound with 0.2 mm enamel-coated copper wire to form an electromagnetic coil. The armature was mounted on the ferromagnetic rail and was free to translate along the longitudinal axis of the capsule.

**Fig. 1.**
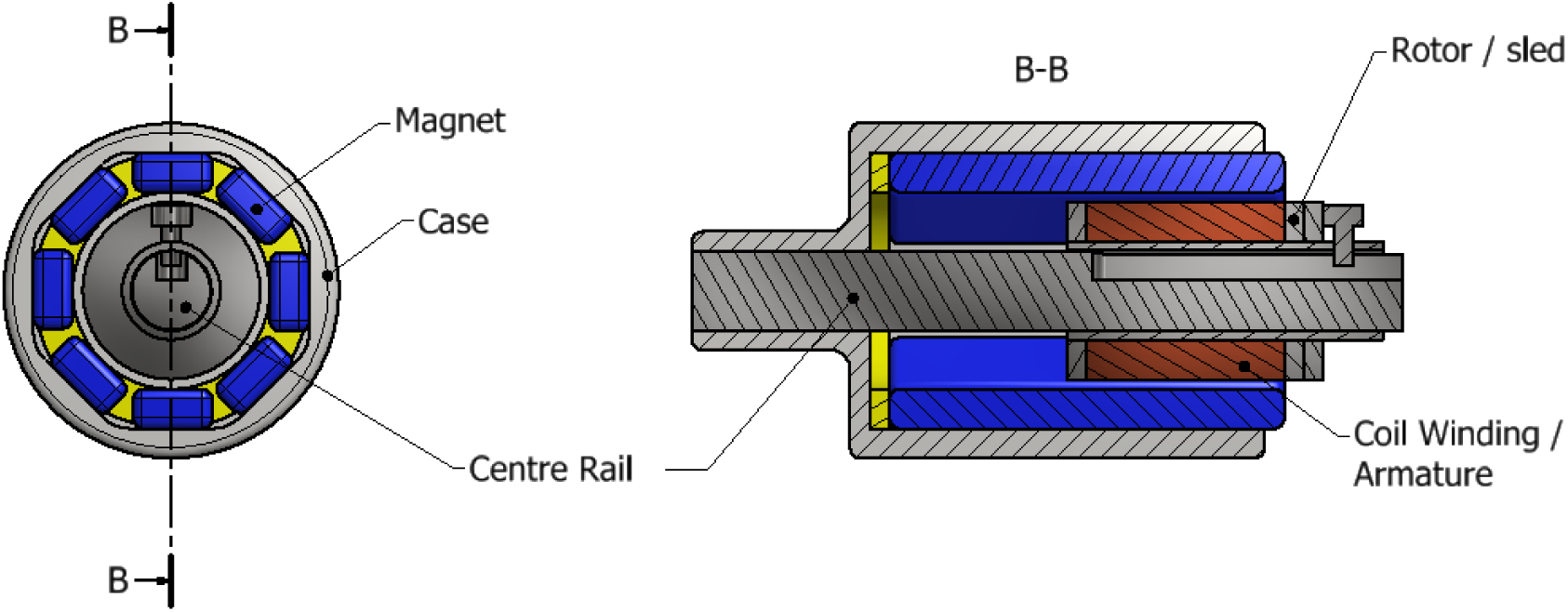
Schematic representation of the electromagnetic impact-actuated locomotion mechanism.

The miniaturised actuator was manufactured using a soft-iron housing, shaft, and end caps with a brass coil bobbin and achieved a substantial reduction in size compared with earlier prototypes. The final actuator measured 12.6 mm in diameter and weighed 11.0 g. For experimental evaluation, the actuator was encapsulated within a custom DLP 3D-printed capsule enclosure with final dimensions of 40.0 mm in length and 15.0 mm in diameter. The total mass of the encapsulated prototype was 12.2 g. This prototype was implemented as a tethered capsule, with external wiring supplying power and control signals to the actuator during benchtop and ex-vivo testing. The tethered configuration enabled rapid optimisation of drive parameters and evaluation of locomotion performance prior to integration of onboard electronics, power supply, imaging, and wireless communication subsystems. When electrical current was applied to the coil winding, interaction between the generated magnetic field and the surrounding permanent magnets produced a Lorentz force, accelerating the armature along the rail.

To achieve locomotion, a current is passed through the coil. As explained earlier, this causes the coil to experience a force. The direction of this force will be in either direction along the axis of the rail (according to the diagram above, either to the left or the right), depending on the direction of the current flowing in the coil. This accelerates the sled-coil assembly. At t_0_, the capsule-mounted sled is stationary. This refers to when t = 0 and t = t_1_ in the series of images below (see Fig. 2). At t_1_, an external magnetic force (F_mag_) is applied but remains insufficient to overcome the frictional resistance (F_fric_). At t_2_, an impact force generated by magnetic attraction exceeds the frictional force (F_impact_ > F_friction_), resulting in rapid sled displacement. At t_3_, the magnetic force is reduced to approximately 10% of its peak value, becoming insufficient to sustain further motion (F_mag_ ≤ F_friction_). At t_4_, the sled reaches a new static equilibrium position within the chamber. Due to Newton’s 3rd law, the force acting on the coil (F_mag_) is also acting on the case, but in the opposite direction. Here, since the case is assumed to be in an intestine, it is assumed that the friction on the case (F_fric_) is greater than or equal to the force acting on the sled. Hence, the case should remain stationary during the acceleration. This is depicted at t = t_1_ in the image. As shown at t = t_2_, the sled will continue to move until it hits the back wall of the case. At this point, the momentum of the sled is transferred to the whole assembly, causing the whole assembly to quickly jerk in the direction that the sled was moving. This impact is what causes a force which is large enough to overcome the friction in the intestine (F_impact_). Newton’s 2^nd^ law states that the force exerted by an object is equal to the rate of change of momentum. During an impact, while there is only a relatively small change in momentum for the case, it happens over a very small space of time. Hence the force which the case exerts on the environment is large enough to overcome friction. This impact principle is similar to how pile drivers work. A small rod is accelerated under the force of gravity until is hits a larger rod which is in contact with the ground. This causes the larger rod to be “hammered” into the ground. Similarly, the case here experiences an impact which drives it through the intestine. The inclusion of the rail (which must be made of a ferromagnetic material - such as mild steel) in the design is an important inventive step as it acts to guide the magnetic field from the magnets to the centre of the rail. As a result, the magnetic field density across the coil is stronger to generate larger impact force to ensure the movement of the capsule at a higher speed overcoming the friction. This is because the magnetic field is arranged more perpendicularly to the coils and the magnetic field magnitude is larger. Without this rail, the magnetic field would quickly dissipate the closer to the centre one got and the direction of the field would also rapidly change, causing the coil to produce less thrust. Therefore, the addition of the rail is an inventive step that significantly increases the available power of the device. As shown at t = t_3_ and t = t_4_, the sled is then drawn back to its starting position inside the case by applying a smaller current in the opposite direction from before. This allows the sled to be drawn back along the rail without causing a large enough impact to overcome environmental friction and hence allowing the capsule to remain stationary. If this sequence were run in reverse (starting on the right side of the case) the case would move in the opposite direction.

**Fig. 2.**
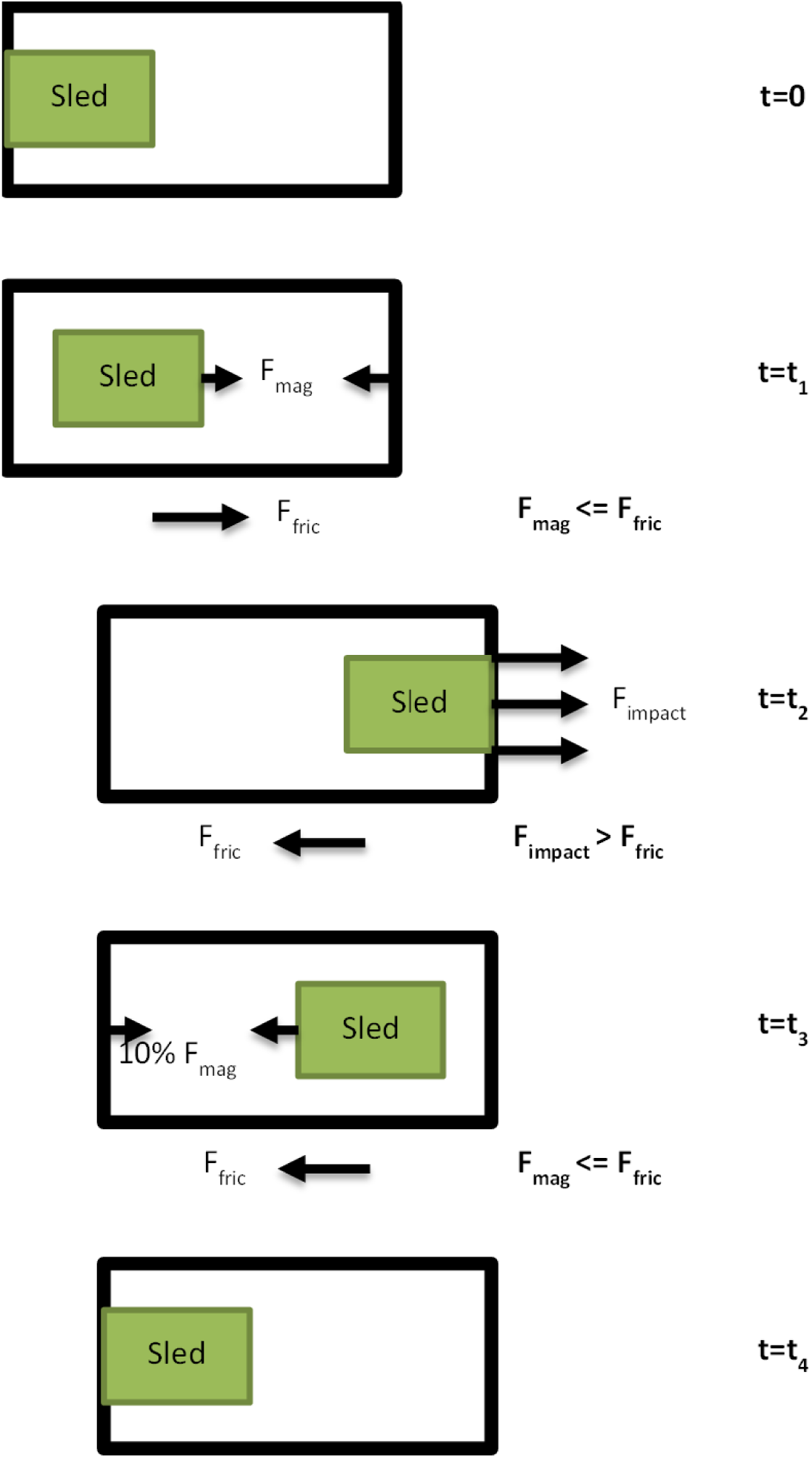
Schematic illustration of magnetic capsule propulsion and sled displacement within the test chamber.

### 2.2 Experimental methods

To ensure safe operation in the intestinal lumen, the helical surface was engineered with a smooth, biocompatible finish to minimise mucosal irritation while maintaining sufficient traction. During the *ex-vivo* trial, the system demonstrated reproducible locomotion in both dry and fluid-filled environments, with measured differences in speed and power consumption reflecting the drag variation across the conditions (the system volume remaining constant throughout). This integration of threaded mechanical propulsion with low-power actuation provided a stable and energy-efficient locomotion strategy, laying the groundwork for future clinical adaptation. The electronic control architecture was based on an Arduino Nano Every microcontroller (Fig. 3), which generated forward and reverse actuation signals using a custom-coded control sequence (Suppl. 1). The actuator was driven *via* a Texas Instruments DRV8837 MOSFET H-bridge IC, selected for its compact size (2.0 mm leadless package), peak current capability of up to 1.8 A, and low internal resistance (typically <0.3 Ω). The microcontroller provided 5 V logic signals, while the solenoid coil was powered by a separate 2.4 V supply, supplied by a two-cell AA battery pack. Transient suppression capacitors (0.1 μF ceramic and 10 μF bulk capacitance) were connected across the actuator supply rail (VM-GND) adjacent to the DRV8837 driver to suppress inductive voltage transients generated during polarity reversal and to improve supply stability during actuation.

**Fig. 3.**
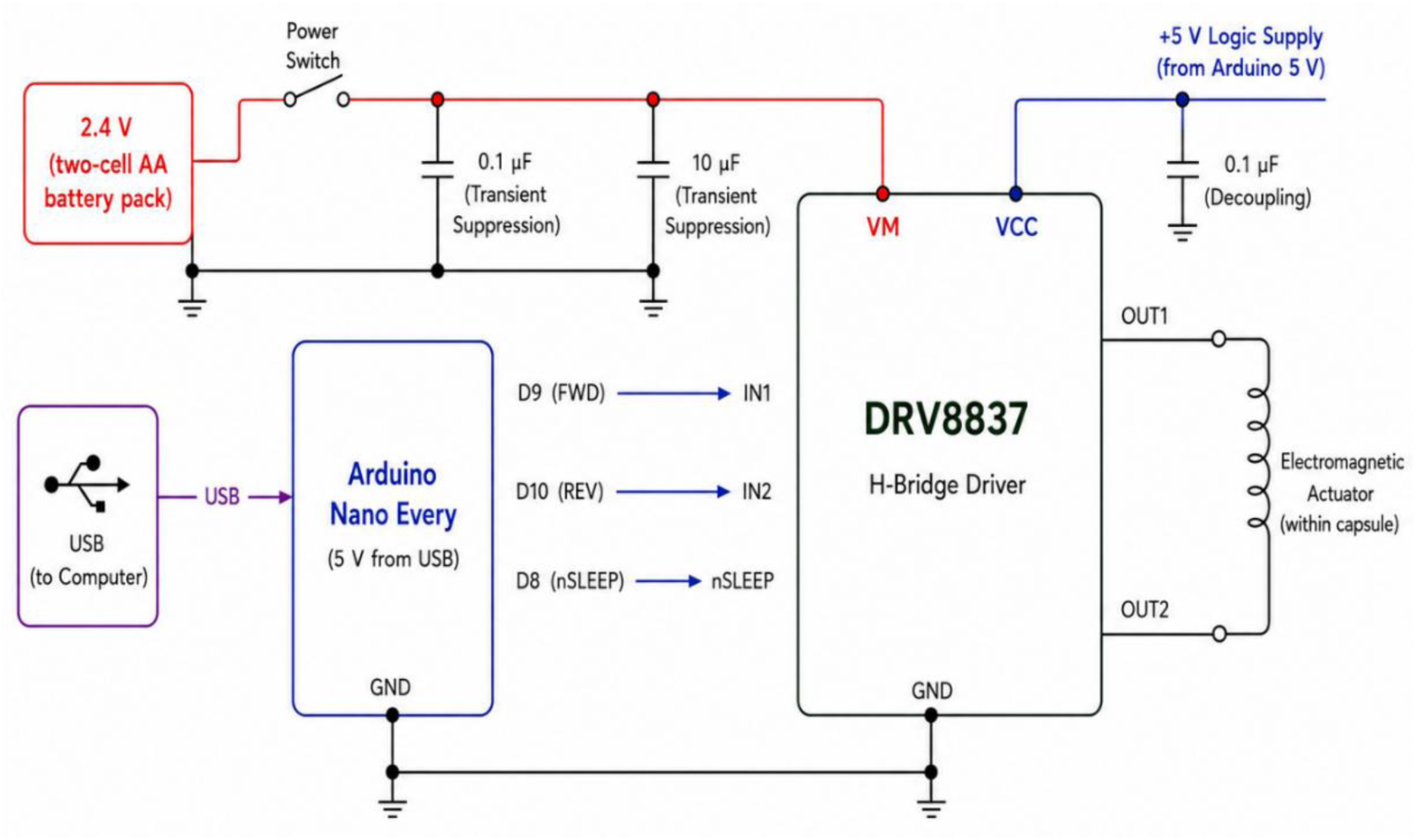
Electronic control architecture of the capsule propulsion system

### 2.3 Bench-test experiment

Prior to biological evaluation, bench-top testing was performed to characterise actuator performance under controlled mechanical conditions. A deformable lumen was simulated using water-filled polyethylene ice-cube bags representing the collapsed intestinal walls with a pressure of ∼90 Pa. The prototype achieved propulsion speeds of up to 8.5 mm/s at 90 Pa, with stalling observed at ∼600 Pa with power consumption of 84 mW. These results validated actuator function and informed parameter selection for *ex-vivo* trials.

### 2.4 *Ex-vivo* Intestinal Model Preparation

We report an active capsule endoscope capable of self-propulsion and directional control within a thawed *ex-vivo* porcine intestine. The intestine was ethically sourced from a certified supplier (Medmeat, UK), compliant with all UK and EEC regulatory standards for animal tissue handling. This approach provides a reproducible, ethically neutral platform for locomotion benchmarking. Segments approximately 35 cm in length were isolated, rinsed with isotonic saline, and assigned to two test conditions:

i. Dry lumen condition: drained segments.
ii. Wet lumen condition: saline-filled segments that better represent *in-vivo* physiological conditions.

The intestinal segment was opened and mounted onto a custom horizontal platform for stability. This configuration allowed controlled capsule deployment and repeatable measurement of propulsion behaviour under both environmental conditions.

### 2.5 *Ex-vivo* experimental Setup and Deployment

Following bench-top validation, the capsule was introduced proximally into each intestinal segment. Experiments were conducted at a controlled room temperature (22-24 °C) on a horizontal rig designed to minimise external friction. Capsule progression was monitored using a high-speed digital video camera, with external tracking markers applied to quantify displacement over time. The capsule endoscope (CE) was evaluated in two different *ex-vivo* porcine bowel environmental conditions: (i) dry lumen and (ii) physiologically simulated wet lumen as shown in Fig. 4 (a and b).

**Fig. 4.**
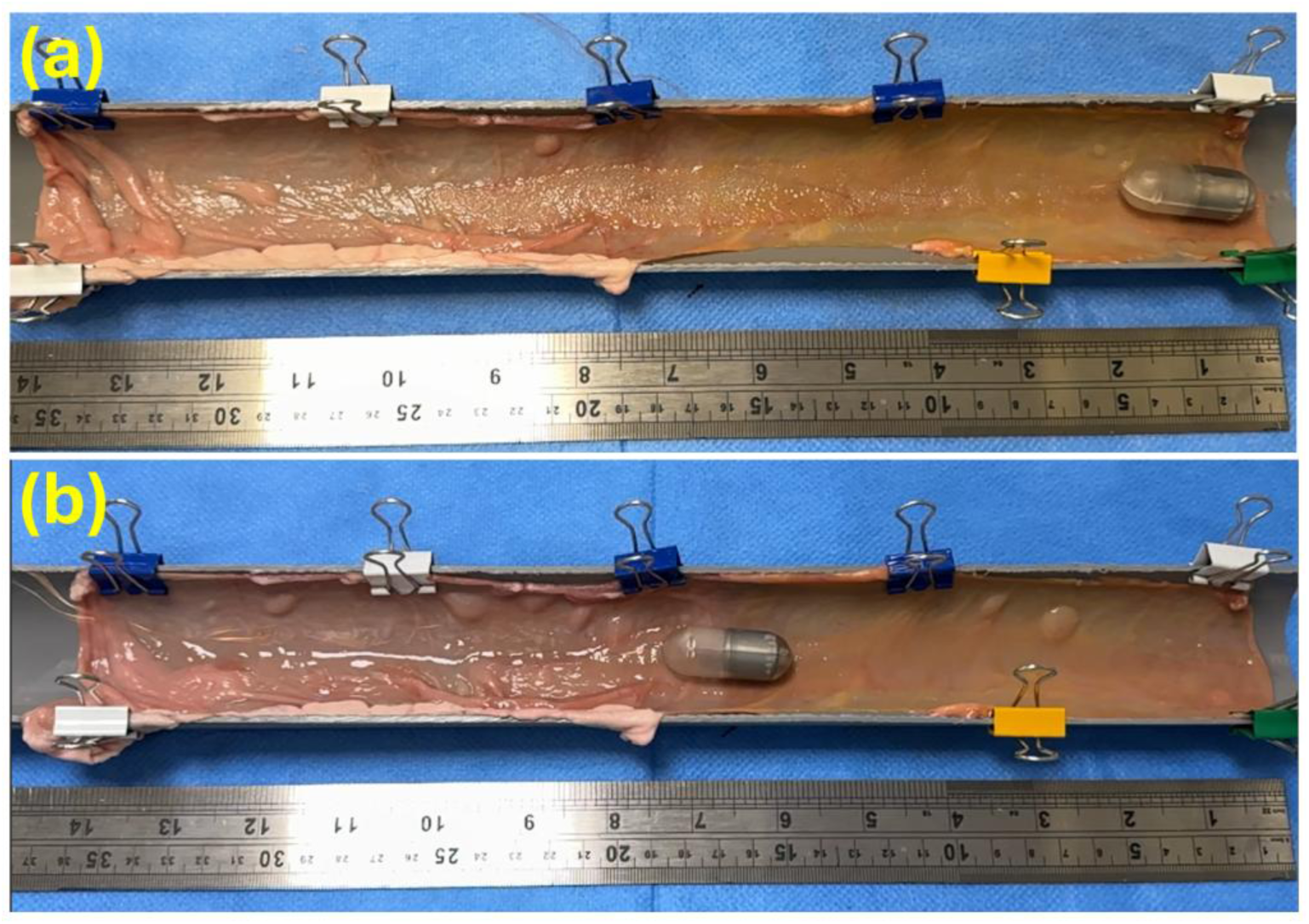
(a) Dry lumen and (b) wet lumen test rigs.

#### Ex-vivo set-ups

**Set-up I:** The bowel was opened along the mesenteric border and placed with the antimesenteric surface uppermost (see Fig. 4a). In this configuration, the capsule moved relatively smoothly along flat surfaces but encountered considerable resistance when traversing irregular folds. Motion was paused in these regions to prevent tether damage, exposing propulsion vulnerability in complex anatomy.

**Set-up II:** The bowel was prepared with saline introduced intermittently throughout each trial to simulate physiological moisture (see Fig. 4b). Saline irrigation reduced surface friction and allowed the capsule to glide more easily. The capsule achieved much faster and more stable transit under wet conditions, although excessive irrigation occasionally created pooling turbulence paradoxically increasing drag and restricted motion. The combined bench-top and *ex-vivo* testing approach enabled comprehensive evaluation of actuator dynamics, energy efficiency, and environmental adaptability, establishing performance benchmarks for future in-vivo trials and clinical translation.

## 3. Results and Discussion

The active capsule endoscope was evaluated under three progressively more realistic conditions - simulated bench, dry lumen, and wet lumen - to assess how environmental factors influence locomotion and energy efficiency. This approach follows the structured evaluative framework outlined in our previous work, combining mechanical, energetic, and translational metrics to provide a comprehensive performance map [24]. Each test involved a fixed 35 cm traversal distance, and the capsule geometry remained constant (length = 40 mm, diameter = 15 mm, volume ∼6.19 cm^3^). Quantitative results are summarized in Table 1. The bench test established the mechanical baseline, with a mean speed of 8.5 mm/s at 84 mW, completing the course in 41 s. Energy expenditure was minimal (9.9 mJ/mm), confirming efficient torque transfer under negligible load.

**Table 1.**
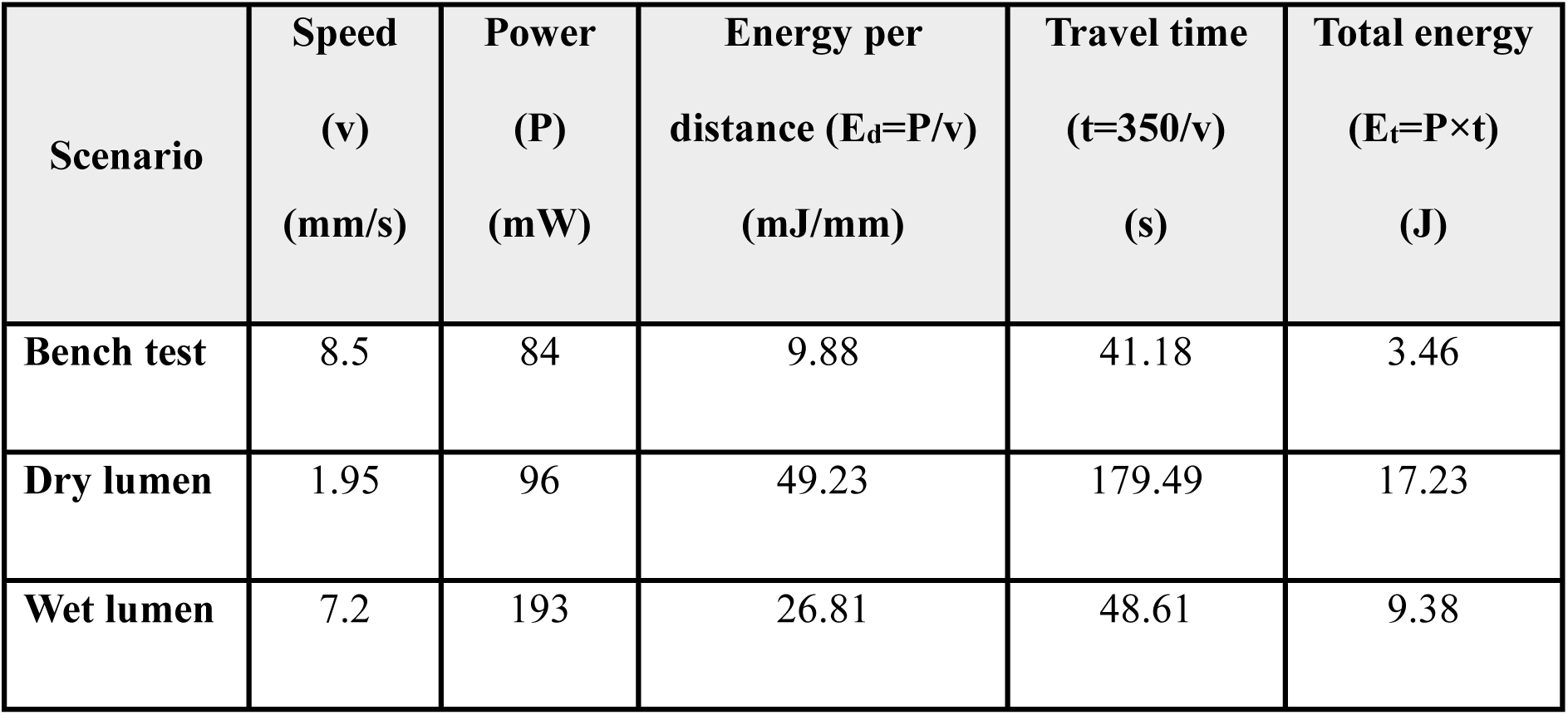
Experimental results (all tests)

| **Scenario** | **Speed**<br>**(v)**<br>**(mm/s)** | **Power**<br>**(P)**<br>**(mW)** | **Energy per**<br>**distance ($E_d=P/v$)**<br>**(mJ/mm)** | **Travel time**<br>**($t=350/v$)**<br>**(s)** | **Total energy**<br>**($E_t=P \times t$)**<br>**(J)** |
| --- | --- | --- | --- | --- | --- |
| **Bench test** | 8.5 | 84 | 9.88 | 41.18 | 3.46 |
| **Dry lumen** | 1.95 | 96 | 49.23 | 179.49 | 17.23 |
| **Wet lumen** | 7.2 | 193 | 26.81 | 48.61 | 9.38 |

This performance defines the upper operational limit for the propulsion mechanism. The results reveal a clear dependence of locomotion behaviour on the environmental medium. In the bench test configuration, the capsule achieved its highest mean velocity of 8.5 mm/s, traversing the 35 cm path in just over 41 s. This condition reflects the intrinsic mechanical capability of the drive mechanism under minimal resistance, establishing the baseline performance limit.

In contrast, during the dry-lumen trials, when the capsule was in direct contact with non-lubricated intestinal tissue, the mean speed dropped precipitously to 1.95 mm/s, extending traversal time to nearly 180 s. Despite a modest increase in average power consumption from 84 mW to 96 mW, translational efficiency collapsed by roughly five-fold, as indicated by the rise in energy-per-distance from 9.9 mJ/mm to 49.2 mJ/mm. This disparity suggests that most of the input power was dissipated as frictional loss rather than productive motion. High static friction and adhesive interaction between the capsule shell and the dry mucosal surface likely induced intermittent stick-slip cycles, characterized by short bursts of movement punctuated by stalls. These cycles waste energy as heat and vibration, providing little net displacement per unit power.

The wet-lumen condition restored much of the capsule’s mobility. Here, lubrication between the capsule surface and the tissue reduced static friction and allowed the capsule to glide more freely, producing a mean speed of 7.2 mm/s that is about 85 % of the bench-top performance. However, this recovery in speed came with a substantial rise in electrical power demand to 193 mW, more than double that of the simulated bench case. The increase in power is attributed to additional viscous drag from the lubricating film and potential internal losses within the motor system due to minor ingress of moisture and increased torque load.

Despite the higher instantaneous power draw, the shorter travel time under wet conditions led to a total energy expenditure of 9.38 J, which was 46 % lower than the dry-lumen case (17.23 J). Expressed as energy cost per distance, the wet-lumen environment required 26.8 mJ/mm, nearly half the expenditure of the dry lumen and only 2.7 times higher than in the ideal bench condition. These results demonstrate that environmental lubrication significantly improves the energy economy of locomotion, even if viscous drag increases total power consumption.

Plotting the data (Fig. 5) reveals a distinct V-shaped trend for both speed (red curve) and power (blue curve) across the three conditions. The minima for both variables occur in the dry-lumen condition, while the bench and wet-lumen conditions cluster at higher values. The pattern reflects a fundamental trade-off between surface friction and fluidic resistance: in the dry lumen, high static friction limits movement despite low viscous load, whereas in the wet lumen, reduced friction permits motion, but this is partially offset by increased hydrodynamic drag. The net effect of lubrication is therefore beneficial, as evidenced by improved speed and reduced energy-per-distance relative to the dry lumen.

**Fig. 5.**
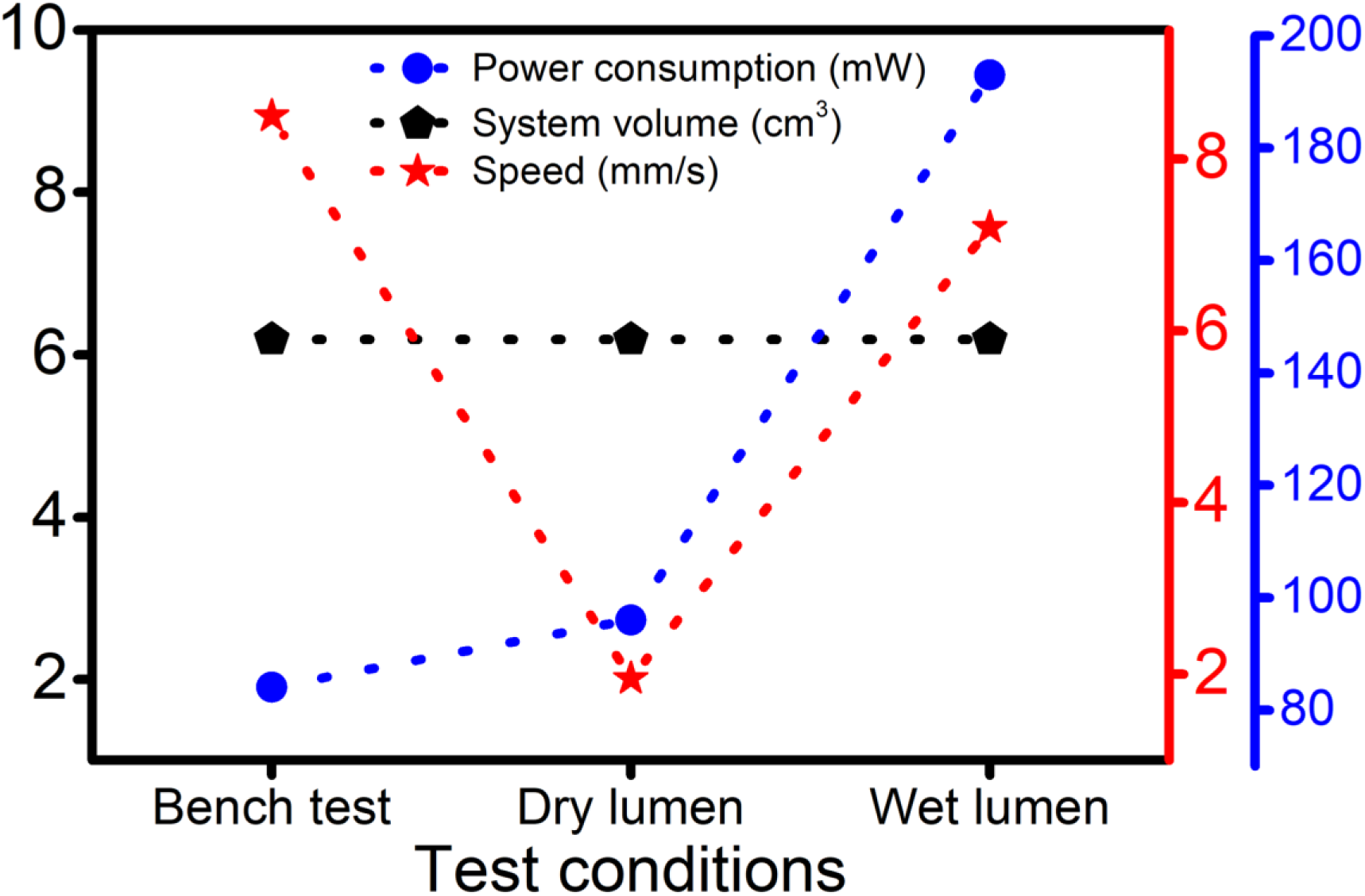
Experimental results of the bench test, dry lumen test, and wet lumen test

### 3.1 Analysis of locomotion efficiency

To quantify propulsion efficiency across test environments, a relative efficiency factor (*η*_*r*_) was defined as the ratio of bench-test energy cost to that of each scenario:

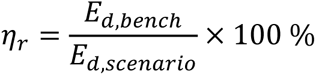

The computed efficiencies were 100 % (bench), 20 % (dry lumen), and 37 % (wet lumen). Hence, wet-lumen operation restored nearly twice the efficiency lost in dry-tissue contact, confirming that lubrication substantially mitigates mechanical losses. The observed decoupling between speed and power in the dry lumen offers important insight into the propulsion physics of capsule robots. Ideally, greater power input should yield proportionally higher speed; however, under high-friction conditions, we encounter the opposite effect in that as power increases, speed drops. This nonlinearity indicates that a critical fraction of the supplied energy is diverted into deforming the tissue surface and overcoming adhesion, not into translational work. Once lubrication re-establishes partial hydrodynamic separation, this coupling is restored, and mechanical output becomes commensurate with electrical input. Taken together, these results establish a clear mechanical hierarchy for propulsion efficiency:

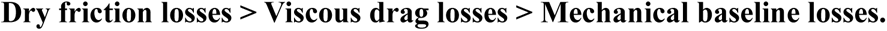

The energy penalty in dry tissue is approximately double that of lubricated conditions and five times that of the ideal bench environment. Overall, the wet-lumen scenario offers the most physiologically relevant performance snapshot, combining near-optimal mobility with a manageable power budget. The data demonstrate that although viscous resistance increases instantaneous power demand, lubrication fundamentally improves propulsive efficiency per unit distance, making it the preferable operational mode for *in-vivo* deployment. The findings confirm that the capsule’s drive system can sustain controlled, forward locomotion in complex biological geometries while maintaining energy consumption within the limits of a practical onboard power source.

Our experimental findings directly validate the evaluative framework established in our earlier study, which proposed a universal scoring and benchmarking methodology for capsule endoscope locomotion systems (CELS) [24]. In that work, we introduced fifteen weighted categories spanning experimental verisimilitude, operational capacity, practicality, and performance to enable standardised, cross-design comparisons. Within that framework, experimental realism (bench, *ex-vivo*, *in-vivo*, and clinical stages) was assigned the highest weighting, reflecting the increasing fidelity of environmental resistance and power consumption observed as prototypes progress from controlled to biological conditions. Our bench and *ex-vivo* trials neatly embody this transition, quantitatively confirming the theoretical predictions of the framework and demonstrating how propulsion efficiency scales inversely with experimental verisimilitude. The bench test corresponds to a verisimilitude level of 0.25, providing the reference baseline against which mechanical losses can be isolated. The *ex-vivo* wet-lumen trials, employing thawed porcine intestine in compliance with UK/EEC standards, represent a verisimilitude level of 0.50, aligning with the *ex-vivo* category defined in the structured framework [24]. The measured decline in relative efficiency from 100% in the bench condition to 37% in the wet-lumen environment fits precisely within the theoretical gradient predicted by the weighted model, which anticipates a 50 - 70% reduction in effective translational efficiency between bench and *ex-vivo* domains. This strong correlation substantiates the analytical validity of the framework and reinforces its applicability to emerging actuation technologies, including tethered and motor-driven designs.

Furthermore, our results confirm the exponential relationships previously derived for performance parameters. As shown in the established framework, capsule top speed (*v*), on-board system volume (*V*), and power consumption (*P*) follow asymptotic trends reflecting the limits of miniaturised actuation. Our measurements across increasing verisimilitude map consistently onto the predicted exponential energy curve, where speed saturates beyond 3 mm/s and efficiency falls sharply as P exceeds 200 mW. Notably, the capsule’s on-board volume of ∼6.19 cm^3^ lies within the transitional range defined in the framework between theoretical ideal and Olympus-benchmark capsule volumes (2.12-8.00 cm^3^), indicating an acceptable design trade-off between power capacity and mechanical integration.

The energy-per-distance metric (*E*_*d*_ = *P*/*v*) derived in this study extends the prior framework by providing an empirical anchor for the power consumption performance category, which had previously been modelled theoretically. The observed values of 9.9 mJ/mm (bench), 49.2 mJ/mm (dry lumen), and 26.8 mJ/mm (wet lumen) confirm the asymptotic behaviour described by the exponential performance functions (equations 2 - 4 in Dinkar et al., 2024 [24]), in which minor increases in load cause disproportionately large energy losses. This alignment demonstrates that the proposed scoring system is not only conceptually robust but experimentally predictive when applied to new capsule designs.

This validation highlights the translational coherence between analytical modelling and practical experimentation. Our results, achieved under *ex-vivo* conditions with biological tissue, reinforce the principle that environmental realism is the dominant determinant of system efficiency, justifying its elevated weighting within the framework. Moreover, by situating our new active locomotion prototype within a previously defined comparative structure, the findings provide a reproducible benchmark for subsequent *in-vivo* testing. In doing so, this study transforms the theoretical evaluation framework from our earlier publication into a validated empirical tool capable of guiding the rational design and optimization of next-generation capsule endoscope locomotion systems.

The performance observed during bench-top and ex-vivo testing can be attributed to the unique architecture of the electromagnetic locomotion system. Unlike many previously reported impact-driven capsule devices in which a permanent magnet acts as the moving mass, the present design employs a movable coil-armature assembly operating within a fixed magnetic field. This arrangement allows the permanent magnets to remain stationary while the lighter armature is accelerated along a guided trajectory. The incorporation of a ferromagnetic rail represents a particularly important design feature. The rail functions both as a mechanical guide and as a magnetic flux concentrator, directing magnetic field lines towards the centre of the actuator. This increases the magnetic field density acting upon the coil winding, thereby increasing the available electromagnetic driving force. Enhanced force generation enables greater momentum transfer during impact events and contributes to improved propulsion efficiency. Furthermore, because the armature moves along a defined rail rather than relying on larger moving magnet assemblies, the design minimises internal friction and improves repeatability of actuation. This simplified architecture facilitates miniaturisation while maintaining adequate propulsion force, an important consideration for future autonomous capsule endoscope development. The experimental findings support this design rationale. Under lubricated conditions the device achieved locomotion speeds approaching those observed during bench-top testing, indicating that the actuator is capable of generating sufficient force to overcome realistic gastrointestinal resistance while operating within clinically acceptable power requirements.

When considered alongside leading locomotion mechanisms reported in the literature (provided in Table 2), the present capsule’s performance occupies a competitive position among the best-documented systems to date. Zhou et al. (2013) demonstrated that spiral capsules driven by an external rotating magnetic field could achieve speeds of 8-10 mm/s in lubricated lumens with excellent controllability; however, these designs required a large external magnet assembly and offered no autonomous capability [25]. In contrast, our prototype reached a comparable speed of 7.2 mm/s in the wet-lumen *ex-vivo* intestine without magnetic actuation, using only a compact internal motor and tethered power. This validates the idea that magnet-free mechanical propulsion can yield equivalent translational performance while simplifying system architecture.

**Table 2.** Comparison table of the existing literature and the present work.

| Study | Actuation / Locomotion Type | Environment Tested | Capsule Size (mm) | Speed (mm/s) | Power Source | Reported Power (mW) | Key Features | Limitations | Ref. |
| --- | --- | --- | --- | --- | --- | --- | --- | --- | --- |
| **Zhou et al. (2013)** | Spiral magnetic rotation (external rotating field) | Bench / fluidic | 13 × 32 | 8 - 10 | External magnetic | ≈150 (field energy) | Efficient in lubricated lumen; controllable rotation | External actuation required; limited autonomy | [25] |
| **Simi et al. (2010)** | Hybrid legged + magnetic | *In-vitro* / *ex-vivo* / *in-vivo* | 14 × 30 (prototype) | 2 - 4 | on board + external field | ≈200 | Hybrid locomotion; manages collapsed regions | Bulky; complex control; scalability | [26] |
| **Valdastri et al. (2009)** | 12-leg lead-screw slot-follower | *Ex-vivo* porcine colon | 11 × 25 | 4 - 6 | On-board DC motor | ≈300 | Strong traction; effective tissue distension | Mechanical complexity; high energy cost | [27] |
| **Formosa et al. (2020)** | Dual motor treaded (Endoculus) | *In-vivo* & *ex-vivo* porcine | 13 × ≈60 (length) | ≈40 | On-board dual DC | >400 | 2-DOF skid-steering; integrated endoscope tools | Large footprint; complex gearing | [28] |
| **Gao et al. (2016)** | Inchworm-like micromotor | Ex-vivo colon | 24 × 33 | 1.2 - 2.6 (≈7.4 cm · min<sup>-1</sup>) | Wireless powered (WPT) | N/A (3.3 V stable) | Tetherless; expandable; safe contact design | Slow locomotion; low torque | [29] |
| **Yim & Sitti (2012)** | Magnetically actuated rolling soft capsule | Bench / synthetic stomach | 10 × 25 | 5 - 8 | External magnetic | External field energy | Soft compliant body; safe surface rolling | Requires precise field control; limited to stomach | [30] |
| **Present work (2025)**<br>Active capsule locomotive system | On-board electric motor | Bench / ex-vivo porcine (dry & wet lumen) | **15 × 40** | **8.5 (bench) / 1.95 (dry lumen) / 7.2 (wet lumen)** | **Tethered DC** | **84 (bench) / 96 (dry lumen) / 193 (wet lumen)** | Efficient wet-lumen locomotion; validated over 35 cm; moderate power; low mechanical loss | Power increases in viscous media; requires miniaturization for ingestible scale | -- |

Simi et al. (2010) advanced a hybrid magnetic-legged capsule that coped with collapsed bowel regions but was limited to speeds of 2-4 mm/s and required intricate coordination between onboard and external components [26]. Our system, operating at almost double that speed in biological media, maintains continuous motion without reliance on legged anchoring cycles, thereby reducing mechanical complexity and potential wear. Similarly, Valdastri et al. (2009) achieved strong traction with a 12-leg lead-screw design operating at around 4-6 mm/s but at a high energetic cost (∼300 mW). In comparison, our capsule approaches their translational output but using less than two-thirds of the power, offering a better speed-to-power ratio and a simpler mechanical envelope [27]. Among the more recent high-power solutions, the dual motor treaded Endoculus developed by Formosa et al. (2020), delivers remarkable manoeuvrability (∼40 mm/s from 400 mW) and colonoscopic functionality, but at the expense of miniaturisation, its body exceeding 60 mm in length [28].

Our prototype is based on a different design philosophy that optimises energy efficiency within swallowable size parameters (15 × 40 mm), achieving biologically useful speeds while consuming moderate power (≤ 193 mW). This positions our capsule nearer the clinically scalable end of the performance spectrum than Endoculus.

Among slow but structurally adaptive systems, Gao et al. (2016) reported an inchworm-type capsule that traversed at only 1.2 - 2.6 mm/s despite wireless power transfer [29]. Although their design ensures safe lumen contact, the extremely low velocity and discontinuous motion contrast sharply with the smoother, continuous propulsion achieved here even in the dry-lumen condition where the capsule-maintained speeds of 1.95 mm/s under severe frictional load. The inchworm paradigm remains advantageous for collapsed segments but unsuitable for efficient endoluminal scanning.

Likewise, the magnetically actuated rolling capsule by Yim and Sitti (2012) could move at 5-8 mm/s demonstrating excellent surface compliance but depended entirely on controlled external fields and performed optimally only within open gastric cavities [30]. Our design offers comparable translational velocity in confined intestinal geometry, confirming that active mechanical propulsion can match magnetic rolling efficiency under *ex-vivo* physiological conditions.

Taken together, these comparisons suggest that the developed system bridges the gap between externally actuated and self-driven capsules. Its speed range (1.95-8.5 mm/s) and power band (84-193 mW) fall squarely between the low-speed inchworm and the high-speed magnetic or spiral classes, achieving an optimal compromise between autonomy, miniaturisation, and energetic economy. In doing so, the device demonstrates that effective propulsion within the gastrointestinal tract need not rely on external fields or oversized actuators, but rather that careful optimization of drive torque, frictional contact, and capsule geometry can deliver near-benchmark performance with practical simplicity. This positions our work as a validated transitional platform between laboratory prototypes and clinically deployable autonomous capsule endoscopes.

However, although the thawed porcine intestine model supplied by Medmeat (UK) conforms to all UK/EEC regulatory standards and provides reproducible geometry and mechanical properties, it does not fully replicate *in-vivo* peristaltic activity or dynamic mucus flow. The absence of peristaltic contraction means that frictional forces are predominantly static, while in living tissue, transient contractions may intermittently assist or oppose motion.

Furthermore, fluid viscosity and mucin content after thawing may differ slightly from physiological conditions, potentially under-representing viscous drag *in-vivo*. Nevertheless, the *ex-vivo* setup represents an ethically sound and controllable intermediate step between benchtop and animal studies, enabling quantitative performance benchmarking without live-tissue variability. Measurement uncertainty in speed (±0.2 mm/s) and power (±5 mW) produces less than ±10 % variation in *E*_*d*_, insufficient to alter the observed trends. Repeated trials confirmed reproducibility within this range.

## 4. Conclusion

This study evaluated an active capsule endoscope’s locomotion across bench, dry lumen, and wet lumen conditions using ex-vivo porcine intestine. Despite identical geometry (volume ∼6.19 cm^3^), performance varied sharply with environment. Speed declined from 8.5 mm/s on the bench to 1.95 mm/s in the dry lumen, recovering to 7.2 mm/s when lubricated. Corresponding power demands rose from 84 mW to 193 mW, while energy cost per distance increased from 9.9 mJ/mm (bench) to 26.8 mJ/mm (wet lumen) and 49.2 mJ/mm (dry lumen). Results show that surface friction dominates energy efficiency, with lubrication restoring mobility and halving total energy expenditure despite higher instantaneous power. The capsule maintained 85 % of its potential speed in wet conditions, confirming robust propulsion in realistic intestinal environments. Overall, the system demonstrates efficient, controllable locomotion and defines clear benchmarks for further optimization. Reducing frictional losses and refining torque control will be key to achieving fully autonomous, energy-efficient navigation for future clinical capsule endoscopy. Future work will focus on refining propulsion technology, extending operational battery life, and validating capsule performance in preclinical *in-vivo* studies. The present study additionally demonstrates the feasibility of a rail-enhanced electromagnetic impact propulsion mechanism for capsule endoscopy applications. By concentrating magnetic flux and improving actuator efficiency, the ferromagnetic rail architecture provides a practical approach for increasing propulsion force without substantially increasing device size or power requirements. This design may represent a promising pathway towards fully autonomous capsule systems capable of active navigation within the gastrointestinal tract. Ultimately, these advances aim to expand capsule endoscopy from a purely diagnostic modality towards a versatile, actively navigated platform capable of targeted interventions and therapeutic applications [31-32].

## Acknowledgments

The authors would like to acknowledge the financial support offered to our Robotic Active Capsule Endoscopy research programme. The work that contributed to enabling this paper was partially funded through the following grants:

1. Project: Miniaturized on-board locomotion system for a capsule endoscope (CE-MOVE); Grant: FAST (Funding at the Speed of Translation), named i4i (Invention for Innovation) by the National Institute for Health and Care Research (NIHR), British Government; Grant code: NIHR206073. This funding was not used for the animal tissue experiments. Instead used to design and fabricate the prototype and complete the bench tests only.
2. Project: Development and evaluation of locomotion mechanisms for robotic active capsule endoscope. Grant: QMI (Queen Mary Innovation) Proof of Concept Award.

The authors would like to express their sincere gratitude to Maddison Limited for providing input into protype fabrication as a partner within the NIHR project, as well as to Ms. Eleanor McAlees, Clinical Trials Research Nurse, Barts and The London School of Medicine and Dentistry, and Mr. Rodrigue Mbuton, Service Manager, Theatre Equipment & Pain Services, Adults, Paediatric Theatres & MEH Theatres, The Royal London Hospital, Barts Health NHS Trust, for their invaluable assistance in the ordering, collection, and preservation of porcine intestinal specimens used in the *ex-vivo* trials. The authors thank Mr. Mish Toszeghi for proofreading the manuscript.

## Declaration of interests

The authors declare no competing interests.

## Data availability statement

The data that support the findings of this study are available upon reasonable request from the authors.

## Author statement

Authors did not use AI and AI-assisted technologies in the writing process.

## Credit authorship contribution statement

Deepak Kumar Dinkar: Methodology / Investigation / Preparing the original draft.

Hasan Shaheed: Conceptualization / Reviewing / Editing.

Kaspar Althoefer: Reviewing / Editing.

Mohamed Adhnan Thaha: Conceptualization / Supervision / Reviewing / Editing.

## Supplementary information file

**Suppl. 1.**
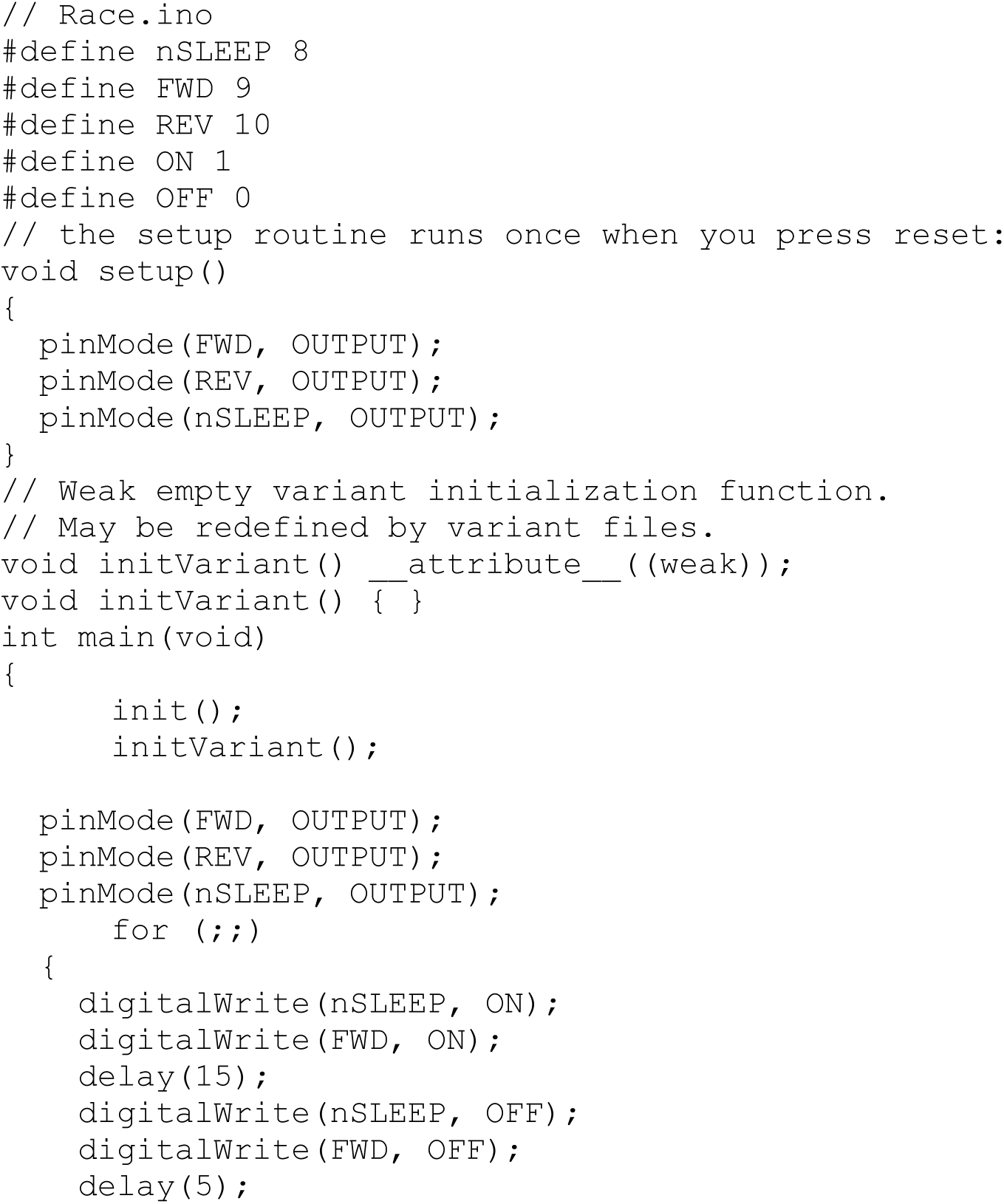
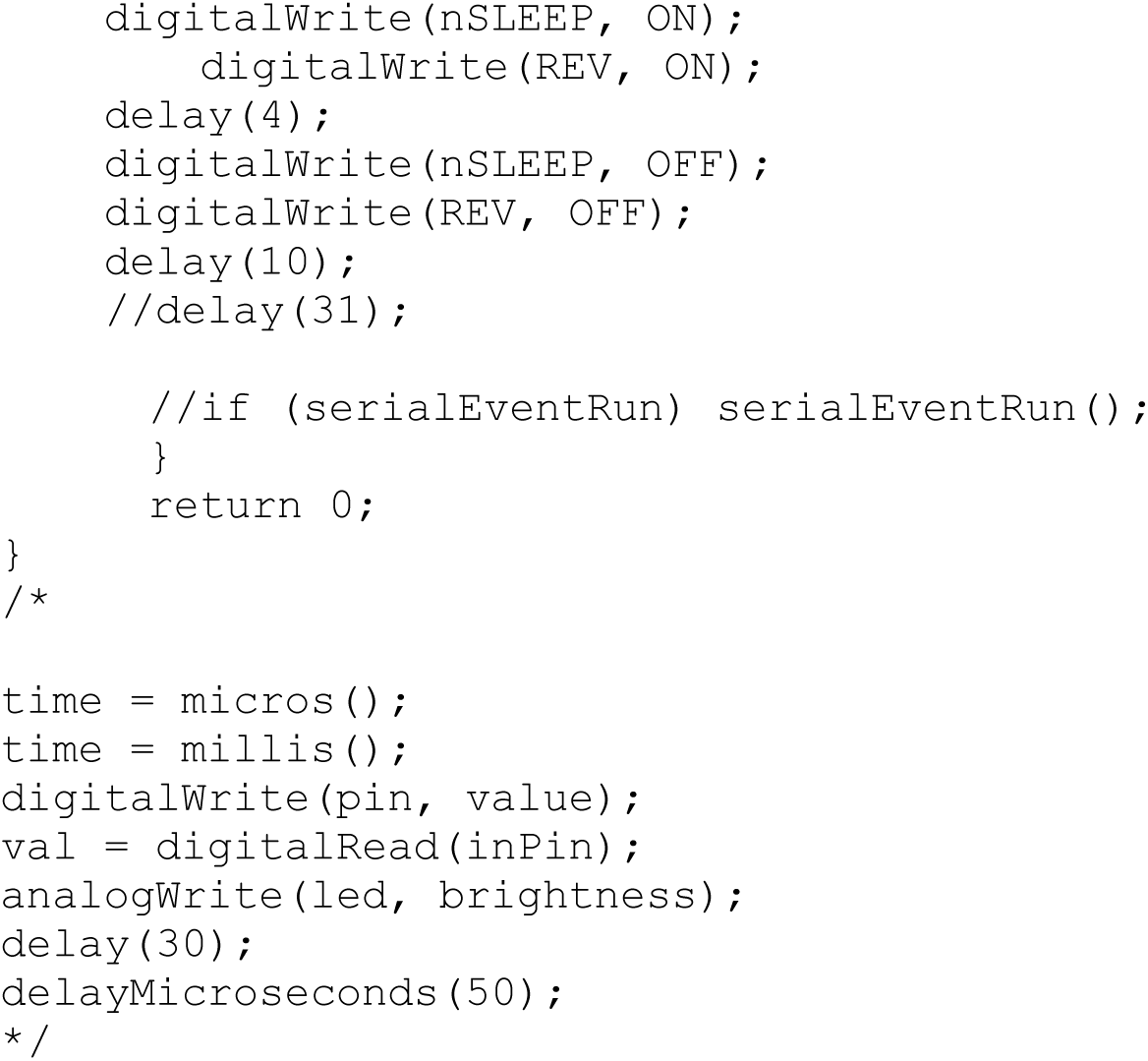
Arduino code used for bidirectionality of the active capsule endoscope.

## Notes

### Competing Interest Statement

The authors have declared no competing interest.

